# The Frizzled-ligand Norrin acts as a tumour suppressor linking oncogenic RAS signalling to p53

**DOI:** 10.1101/666925

**Authors:** Gabriele Sulli, Errico D’Elia, M. Cristina Moroni

## Abstract

Tumour suppressor genes are frequently affected by somatic alterations in cancer, and the impairment of their normal function provides a strong contribution to tumourigenesis. Short-hairpin (sh) RNA library screens have been employed as powerful genetic tools to uncover important new players in human cancer ^1–5^. To identify potential novel tumour suppressor genes acting in the p53 pathway, we performed an shRNA screen using a cell-based model in which only a single additional genetic event disrupting the p53 pathway is required to obtain *in vitro* transformation. By using this approach, we report here on the identification of the Frizzled-ligand Norrin (Norrie disease protein) as a candidate tumour suppressor. Inhibition of Norrin expression promotes anchorage-independent growth, confers a strong growth advantage to cells and causes a reduction in p53 protein levels. Conversely, recombinant human Norrin increases p53 levels in a β-catenin dependent fashion. Interestingly, Norrin expression is stimulated by oncogenic H-RAS and BRAF, suggesting that Norrin is part of an early fail-safe mechanism to suppress transformation, and that mutation or down regulation of Norrin could contribute to tumour progression. Indeed, we found that Norrin expression is significantly decreased in melanoma, breast, prostate and ovarian cancer. These findings support the existence of a novel autocrine/paracrine feedback loop that constrains tumourigenesis, in which the crosstalk between the RAS and β–catenin pathways play an unanticipated role.

## Introduction

The development of in vitro step-wise transformation assays using normal diploid human cells of different origins has provided important insights into the molecular mechanisms leading to cancer ^6,7^. These assays in combination with the availability of short-hairpin (sh)RNA libraries enable loss-of-function screenings in mammalian cells^8^.

To identify novel candidate tumour suppressor genes with a potential function in the p53 pathway, we performed a retroviral shRNA library screen in a human cell model prone for transformation (Fig. 1a). We used the human diploid lung fibroblast cell line, TIG3 as a model system for transformation. These cells are transformed when expressing the human telomerase catalytic subunit (hTERT), an shRNA targeting pRB, an shRNA targeting p53, H-RASV12 and the SV40 small t-antigen (ST). In the cell system used for the screen we omitted the shRNA targeting p53, and the transformation prone TIG3 cells therefore expressed TERT, shRNA pRB, ST and H-RASV12. These cells were infected with a retroviral shRNA library targeting approximately 8000 human genes ^1^ and selected for anchorage-independent growth (Fig. 1a). The shRNA inserts from the recovered colonies were then amplified by PCR, and re-cloned for sequencing and validation.

**Figure 1.**
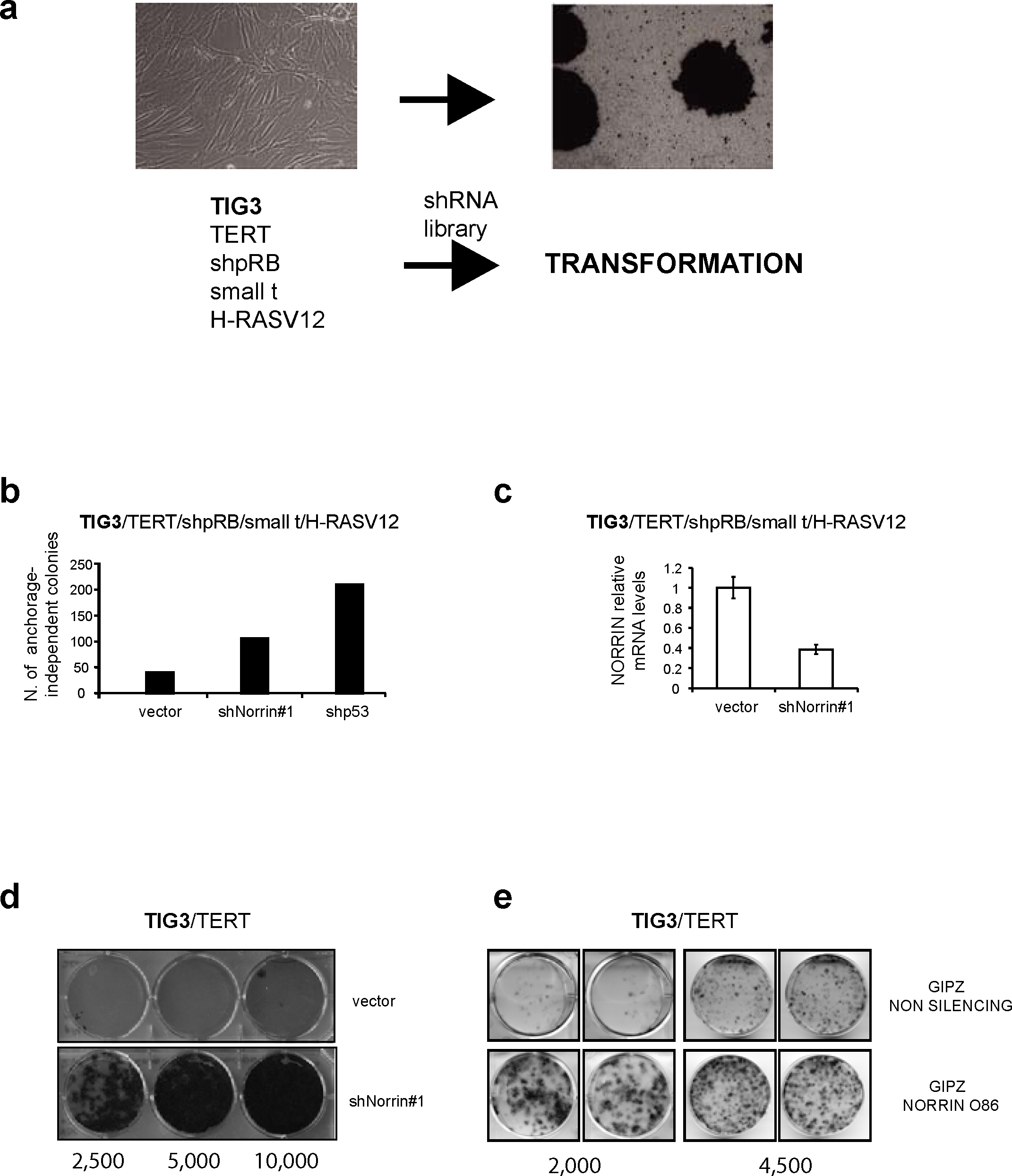
Identification and validation of Norrin in a screening for candidate tumor suppressor genes. **a.** Outline of the transformation screen for the identification of putative tumour suppressor genes acting in the p53 pathway. **b.** Inhibition of Norrin expression leads to anchorage-independent growth of transformation-prone TIG3 cells. The TIG3 transformation-prone cells were infected either with the empty pRetroSuper (pRS), the pRS encoding Norrin shRNA (shNorrin#1) identified from the screen, or pRSp53 as a positive control. The infected cells were cultured in soft agar for 3 weeks. The number of anchorage-independent colonies of a representative experiment is shown. **c.** RT-Q PCR of cells in **b,** with Norrin specific primers. Error bars represent the mean +/-SD for a representative experiment performed in triplicate. **d., e**. Norrin inhibition confers increased ability to grow at low cell density to human diploid fibroblasts. TIG3-T cells were infected either with (d) pRS (vector), or pRSNorrin #1, or (e) pGIPZ Non silencing, or pGIPZNorrin 086. After the end of selection the indicated cell numbers were plated, and stained two weeks later.

## Results

The quality of the screen was underscored by the fact that we identified an shRNA targeting p53 in our screen (data not shown). Among other shRNAs, we identified one targeting Norrin (shNorrin#1). Norrin is a cystein-knot protein whose mutation is associated with the Norrie disease, a syndrome characterized by congenital blindness, deafness and mental retardation ^9–11^. To validate the screen and Norrin as a putative tumour suppressor gene, we re-infected the cells used in the original screen with shRNANorrin#1 and determined the number of anchorage-independent colonies. The cells were also infected with an empty retrovirus as a negative control and a retrovirus expressing an shRNA to p53 as a positive control. As shown in Figure 1b, shNorrin#1 was indeed able to confer anchorage-independent growth to the transformation prone diploid fibroblasts. By RT-QPCR, we confirmed that the endogenous Norrin mRNA level was specifically reduced by shNorrin#1 in the same experiment (Fig. 1c).

Since loss of p53 bypasses proliferation arrest induced in diploid cells grown at low cell density ^1^ we tested whether the inhibition of Norrin expression could confer a similar phenotype. Thus, we infected TIG3-TERT (TIG3-T) cells with two different vectors expressing shRNAs to Norrin and the appropriate controls, and indeed, the inhibition of Norrin expression allowed cells to grow more efficiently when seeded at low density (Fig. 1d, e; Supplementary Figure 1a, b).

Next, we asked whether acute Norrin depletion could affect the p53 pathway. We stably expressed two shRNAs targeting Norrin in TIG3-T cells and determined the expression levels of p53 by immunoblot. Downregulation of Norrin expression led to a substantial reduction in p53 protein levels (Fig. 2a), without affecting p53 mRNA levels (Fig. 2b).

**Figure 2.**
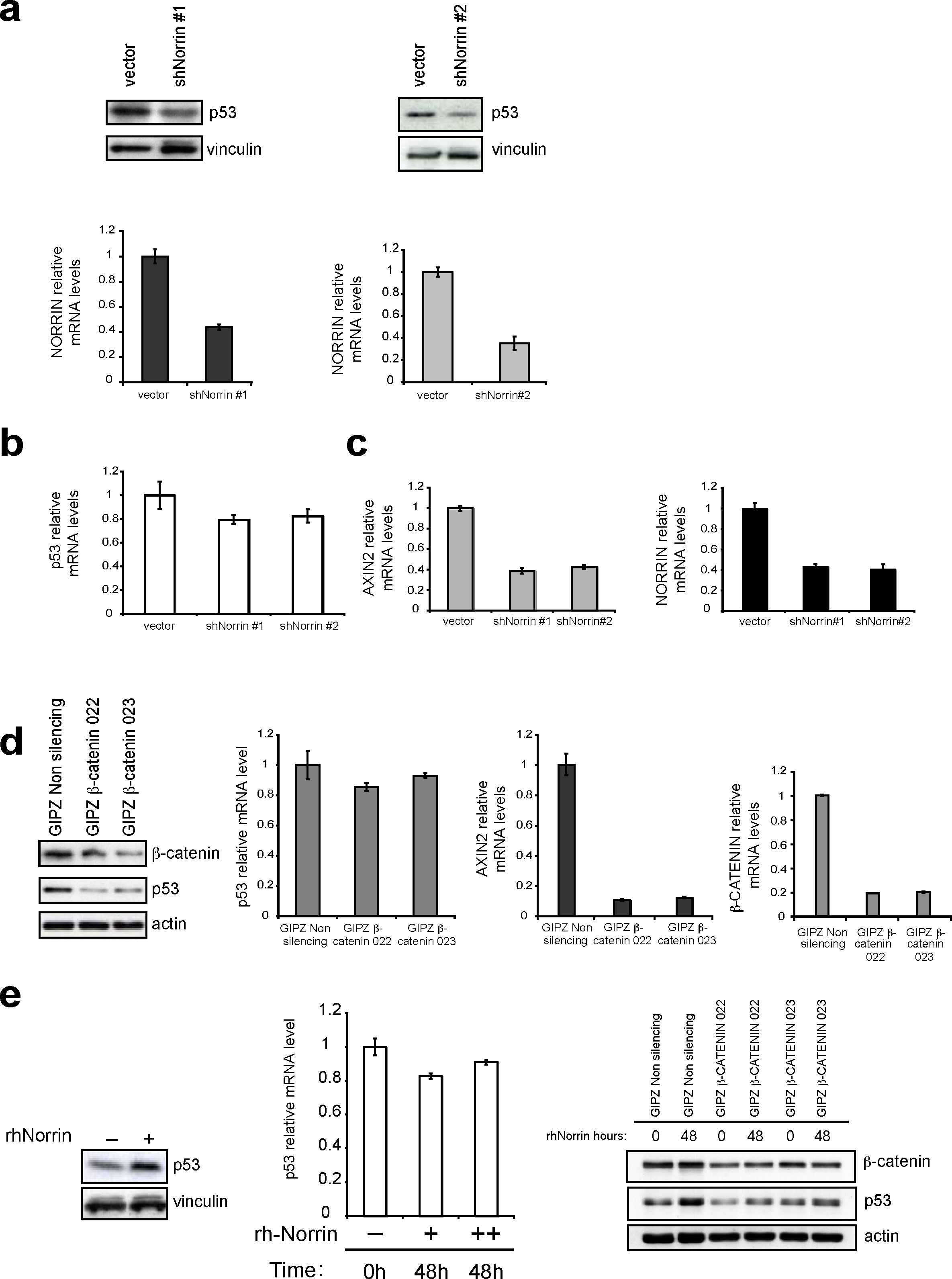
Norrin regulates p53 protein levels through β-catenin. **a.** Inhibition of Norrin expression leads to decreased p53 protein levels. Top panels: Immunoblot analysis of lysates from TIG3-T cells infected with pRSuper, shNorrin#1 or shNorrin#2, showing the p53 protein. Lower panels: RT-Q PCR analysis of Norrin mRNA expression. **b.** p53 mRNA levels are not altered by Norrin knock down. RT-QPCR for p53 mRNA levels, using mRNA from the experiments shown in panel a. **c.** Norrin depletion leads to decreased expression of the WNT/β-catenin target gene *AXIN2*. Left panel: RT-QPCR of *AXIN2* mRNA, using mRNA harvested from TIG3-T cells infected with pRS empty vector, shNorrin#1, or shNorrin#2. Right panel: RT-Q PCR for Norrin mRNA from the same experiment. **d.** β-catenin knock down decreases p53 protein levels without affecting p53 mRNA levels. Left panel: Lysates from TIG3-T cells infected with the indicated pGIPZ interfering constructs were immunoblotted against specific antibodies for the indicated proteins. Right panels: RT-QPCR analysis using mRNA prepared from the cells shown in left panel with primers specific for p53, AXIN2 or β-catenin. **e.** Recombinant human Norrin (rhNorrin) increased p53 protein levels in a β-catenin dependent manner. Left panel: TIG3-T cells were incubated with or without rhNorrin (60 ng/ml) for 48 hours. Cell extracts were immunoblotted for p53 or Vinculin (loading control). Middle panel: RT-QPCR with p53 primers after treatment with 40ng/ml (+) or 80ng/ml (++) of Norrin for 48 hours. Right panel: TIG3-T cells, infected with the indicated constructs, were incubated with rhNorrin (60 ng/ml) for 48h. Cell extracts were immunoblotted for p53, β-catenin or actin (loading control).

Interestingly, Norrin was proposed to be a novel ligand for the Frizzled-LRP receptor complexes ^12,13^. Norrin is a secreted protein, which acts through the Frizzled 4 (FZD4) receptor to activate the WNT/β-catenin pathway ^12,13^. Consistent with this, inhibition of Norrin expression led to the down regulation of the β-catenin target gene *AXIN2* (Fig. 2c, Supplementary Fig. S2a, b).

Increased β-catenin activity was previously shown to increase p53 levels and the activation of a p53-mediated anti-proliferative response ^14,15^. Since Norrin increases the expression of β-catenin target genes, the β-catenin pathway may link Norrin to p53. Consistent with this, inhibition of β-catenin led to reduced p53 protein levels, while p53 mRNA levels were not affected (Fig. 2d).

In agreement with previous studies ^12,13^ we also confirmed Norrin as an activator of the WNT/β-catenin pathway. The treatment of TIG3-T cells with recombinant human Norrin (rhNorrin) led to increased expression of *AXIN2* mRNA (Supplementary Fig. 2c). Furthermore, treatment with rhNorrin of the U2OS human osteosarcoma cell line, led to β-catenin stabilization (Supplementary Fig. 2d).

Next, to complement our results showing that inhibition of Norrin and β-catenin expression reduces p53 protein levels, we asked whether exposure of TIG3-T cells to rhNorrin results in increased p53 protein levels. As shown in Figure 2e, the treatment of TIG3-T cells with rhNorrin indeed triggers p53 accumulation, without leading to a significant increase in p53 mRNA levels. Interestingly, the accumulation of p53 induced by rhNorrin is decreased in cells where β-catenin is down regulated (Fig. 2e). Taken together these results suggest that increased Norrin levels result in the accumulation of p53 through the activation of β-catenin.

Moreover, treatment of TIG3-T cells with rhNorrin induces a p53-dependent inhibition of cell proliferation (Supplementary Fig. 3), suggesting that Norrin, through the activation of β-catenin, triggers a p53-mediated anti-proliferative response.

**Figure 3.**
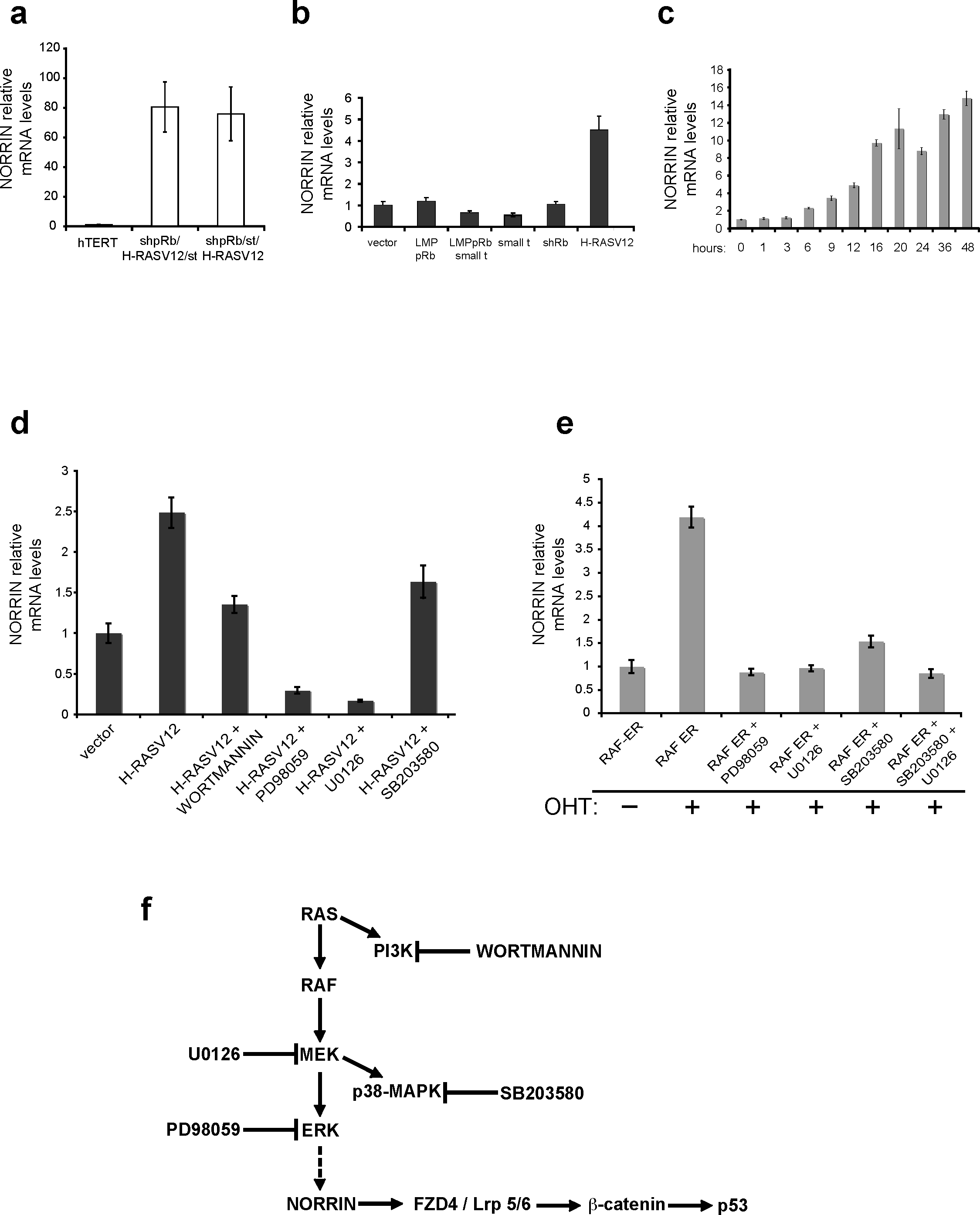
Norrin expression is induced by the RAS-RAF pathway. **a.** Norrin mRNA is expressed at high levels in transformation prone TIG3-T cells (pRS RB, Small T antigene (ST) and H-RASV12) as compared to TIG3-T control cells. Norrin mRNA levels were measured by RT-QPCR. **b.** Oncogenic H-RASV12 induces Norrin expression. TIG3-T cells were infected either with LMP pRB, pBabeST, shRB or pBabe H-RASV12. Norrin mRNA levels were determined by RT-QPCR. **c.** Norrin mRNA is induced by BRAF. mRNA levels of Norrin mRNA determined by RT-QPCR of mRNA harvested from TIG3-T cells expressing BRAF-ER. BRAF activity was induced by OHT treatment for the indicated times (h= hours). **d.** The MEK/ERK pathway activity is required for H-RASV12 induced expression of Norrin. RT-QPCR analysis of Norrin mRNA levels from TIG3-T cells infected with H-RASV12 expressing retrovirus and treated with the indicated inhibitors for 24 hours. U0126: MEK inhibitor; PD98059: ERK inhibitor; Wortmannin: PI3K inhibitor; SB203580: p38/MAPK inhibitor. **e.** The MEK-ERK pathway activity is required for the induction of Norrin expression by BRAF. RT-QPCR analysis of Norrin mRNA from TIG3-T BRAF-ER cells treated for 24h with OHT plus the indicated inhibitor. **f.** Illustration of the Norrin-p53 pathway with the inhibitors used in panels d,e indicated. Norrin provides a novel molecular link between RAS-RAF-MEK-ERK oncogenic signalling and p53, through activation of the WNT/β-catenin pathway.

When cells are affected by oncogenic stimuli that lead to inappropriate proliferative signals, tumour suppressor genes are frequently activated ^16,17^. Thus, we asked whether Norrin expression is increased in response to oncogene activation. We found that Norrin is expressed at low levels in TIG3-T cells, while it is dramatically increased in the transformation-prone cells used in the screens (Fig. 3a). Next, we investigated which of the oncogenic stimuli present in the transformation prone cells could lead to increased levels of Norrin. As shown in Figure 3b, H-RASV12 expression led to a significant increase in Norrin levels (Fig. 3b).

RAS activates multiple signalling cascades including AKT, RALGDS, RAC, MAPK and ERK ^18^. To further explore the mechanism by which RAS signalling induces Norrin expression, we investigated whether activation of RAF could induce Norrin mRNA as efficiently as oncogenic RAS. Therefore, we used TIG3-T cells expressing a BRAF-ER fusion protein, which is activated by the addition of 4-hydroxy tamoxifen (OHT). Indeed, the activation of RAF led to a strong increase of Norrin mRNA (Fig. 3c).

To further investigate the cascade involved in H-RASV12 - induced increase in Norrin expression, we treated cells expressing H-RASV12 or BRAF-ER with inhibitors of PI3K (Wortmannin), MEK/ERK (U0126, PD98059) and p38/MAPK (SB203580) kinases. Norrin induction by H-RASV12 and BRAF was efficiently blocked by U0126 and PD98059 alone, whereas Wortmannin and SB203580 had partial inhibitory effects (Fig. 3d). When SB203580 was combined with U0126, Norrin induction by BRAF was at levels similar to U0126 treatment alone (Fig. 3e). Taken together, these data suggest that RASV12 induces the expression of Norrin through the RAF-MEK-ERK pathway (Fig. 3f).

Since our results demonstrated that Norrin acts as a tumour suppressor in a transformation assay, we analyzed the expression levels of Norrin in cell lines and primary tumours. Melanomas are frequently affected by oncogenic mutations in *RAS* and *BRAF* ^19^. Thus, we analysed whether Norrin expression is decreased in primary and metastatic melanoma cell lines. Indeed, Norrin expression level is decreased in metastatic melanoma cells (Fig. 4a, b, c); in addition, analyses of patient - matched primary and metastatic melanoma cell lines further confirmed that Norrin is strongly downregulated in metastasis (Fig. 4 b, c). To obtain further information on Norrin expression in other cancer types, we searched the Oncomine database ^19^. Norrin expression is significantly decreased in aggressive breast tumours as compared to the more benign stages (Fig. 4 d, e). In addition, Norrin expression inversely correlates with tumour grade in prostate and endometrial cancer (Fig. 4f, g), and is significantly down regulated in ovarian and prostate tumours when compared to normal ovarian and prostate tissues (Supplementary Fig. 4 a, b).

**Figure 4.**
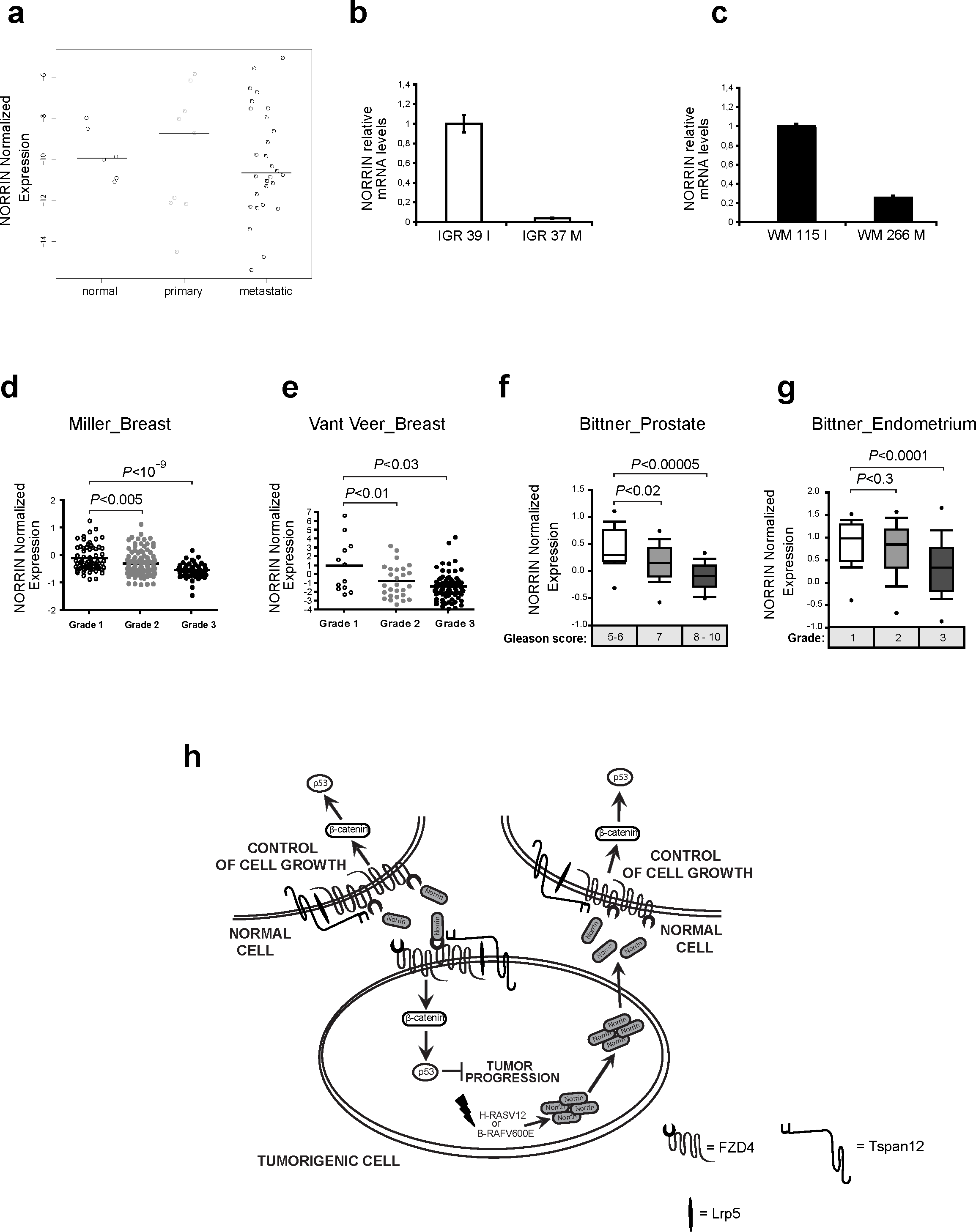
Norrin expression is down regulated in human tumours. **a.** Norrin expression is significantly decreased in melanomas. RT-QPCR analysis of Norrin mRNA from normal melanocytes, primary melanoma cell lines, or metastastic melanoma cell lines. Norrin mRNA levels are expressed relative to RPPo. Bars indicate medians. Norrin mRNA was undetectable in MEL HO and WM1552 primary melanoma cell lines and in CHL1 metastatic melanoma cell lines. Details on the specific cell lines are provided in the Supplementary methods section. **b, c.** Norrin expression is strongly decreased in two metastatic melanoma cell lines versus their matched primary melanoma cell lines. RT-QPCR analysis of Norrin mRNA levels from cell lines derived from primary or metastatic melanoma from individual patients: IGR 39I, IGR37 M, or WM115 I, WM266 (I = Primary, M= metastatic). **d, e.** Down-regulation of Norrin expression during breast cancer progression. Norrin mRNA levels show inverse correlation to the tumour stage, in two different studies from the Oncomine database: Miller et al (**d.**), and Vant’Veer et al (**e.**) P - values of T-tests are provided. **f, g.** In prostate and endometrial tumour progression, Norrin is progressively down regulated. Data from the Oncomine database, P values of T-test are shown. **h**. Model for the role of Norrin in tumourigenesis. Oncogenic activation of RAS or BRAF leads to increased Norrin expression. Norrin activates β-catenin, through an autocrine-paracrine mechanism, and leads to increased p53 levels. Stabilization of p53 by Norrin therefore provides an antiproliferative signal in response to oncogenes. In normal cells, not affected by oncogenic alterations, basal levels of Norrin signalling modulate normal growth.

## Discussion

The WNT/β-catenin signalling pathway is frequently activated in human cancers ^20^. However, as already shown for other major oncogenic signalling pathways, most notably the RAS-RAF pathway, oncogene activation, in this case β-catenin, promotes accumulation of p53 thus leading to growth arrest ^14,15^.

Here we have shown that Norrin can act as a tumour suppressor gene. Indeed, depletion of Norrin expression by shRNA can substitute for the inactivation of p53 in cellular transformation assays.

Increase of Norrin levels induced by oncogene activation activate β-catenin, most likely through an autocrine/paracrine pathway involving the binding of Norrin to the Frizzled receptors, FZD4/LRP5 and the TSPAN12 co-receptor ^21,22^. Subsequently Norrin stimulates a β-catenin dependent accumulation of p53, which in turn triggers a p53-mediated anti-proliferative response (Fig. 4h). Therefore, Norrin is part of a fail-safe mechanism preventing tumour-induction elicited by oncogenic RAS-signalling through the RAF-ERK cascade.

Our findings suggest that Norrin, a non-conventional WNT ligand, could be a previously unappreciated secreted tumour suppressor. Indeed, Norrin expression is triggered by aberrant oncogenic stimuli, positively regulates p53 stability and is downregulated in multiple tumour types and during cancer progression.

Since Norrin acts as a WNT ligand, which is induced by the RAS-RAF pathway, we provide a novel link between the RAS and WNT/β-catenin pathways. Further investigations are required to elucidate a potential role for Norrin in tumourigenesis.

## METHODS

### Plasmids

The following plasmids were used: pWZL-Neo Ecotropic receptor ^23^, pBABE-Hygro hTERT ^6^, pBABE-Puro H-RasV12 ^23^, pBABE-Neo H-RasV12 ^23^, pBABE-Zeo small T ^24^, pRETROSUPER shp53 ^25^, LMP p53 (see supplementary methods), pRETROSUPER shpRb ^25^, GIPZ non silencing, GIPZ Norrin 084/086, and GIPZ -β-catenin 022/023 (Open Biosystems).

### Cell lines and Tissue culture

The human diploid lung embryonic fibroblast cell line TIG3 ^26^, was maintained in DMEM supplemented with 10% (v/v) FBS and 2 mM Glutamine. TIG3 BRAF-ER cells were obtained from Karl Agger.

Ecotropic or amphotropic retoviral supernatants were produced by transfection of Phoenix ecotropic or amphotropic packaging cells ^27^ with the calcium phosphate precipitation technique. At 48 h post-transfection, viral supernatants were collected, filtered, supplemented with 5 μg/ml polybrene and used for infection of the TIG3 cells. Drug selections in TIG3 cells were performed with 1μg/ml puromycin, 50 μg/ml hygromycin, 100μg/ml G418, and 30μg/ml Zeocin.

Cells infected with pBABE-Neo H-RASV12 were treated for 24 hours with wortmannin (Calbiochem) 100nM, PD98059 (Calbiochem) 50μM, U0126 (Calbiochem) 10μM, SB203580 (Calbiochem) 20μM. Cells infected with pBABE-PURO B-RAF-ER were cultivated in the presence of OHT (600nM) and PD98059 (Calbiochem) 50μM, U0126 (Calbiochem) 10μM, SB203580 (Calbiochem) 20μM for 24 hours.

### Anchorage independent growth

Approximately 100,000 cells were plated in 0.45% low melting temperature agarose (Sea Plaque GTG agarose)/growth media onto 6-well dishes with a 0.9% agarose underlay. Colonies were counted from triplicate wells after three weeks, and the total number is shown.

### Colony-formation assays

TIG3-T cells were infected with the indicated vectors, and plated after antibiotic selection at the indicated cell numbers in 6 cm or 10 cm dishes and stained with Crystal violet after 2 weeks.

### Growth curves

TIG3 cells were infected with the indicated retroviral vectors, and after selection 30,000 cells were plated in 6 well plates (time = 0 days). Every 3 days, cells were counted and 30,000 cells were replated. Total cell amounts in all growth curves were displayed as cumulative over time, and calculated as in ^25^.

### shRNA sequences

The targeting sequences of the shRNAs present in the retroviral plasmids are listed in Supplementary methods.

### Data mining

Information regarding data mining is given in supplementary methods.

### Additional materials and methods

Additional materials and methods are described in the supplementary method section.

## Acknowledgments

We thank Kristian Helin for his continuous support and comments on the manuscript. We thank Pier Giuseppe Pelicci for support and comments on the manuscript. We thank René Bernards for the NKI shRNA library, Karl Agger for TIG3 BRAF-ER cells, Paul Cloos and Giovanni d’Ario for help with statistical analysis, and Luisa Lanfrancone and Meenhard Herlyn for the primary and metastatic melanoma cell lines. This work was supported by the Italian Association for Cancer Research (AIRC), the Danish Cancer Society, and the European Framework 6 programme to INTACT.

