## Supplementary info for "The Frizzled-ligand Norrin acts as a tumour suppressor linking oncogenic RAS signalling to p53"

### SUPPLEMENTARY METHODS

#### Cell lines

Primary melanocytes were maintained in McCoy's 5A with glutamax, 2% FBS, Cholera Toxin 20 pM, Hydrocortisone 0.5 μg/ml, Transferrin 5g/ml, TPA 16nM, SCF 10ng/ml, bFGF 1ng/ml, Endothelin-1 10nM

Primary melanoma cell lines were cultured as indicated below:
IGR39 I: DMEM, 10% FBS, 2 mM Gln.
WM 115 I: EMEM, 0.1mM NEAA, 1 mM NaP, 10% (v/v) FBS, 2 mM Gln.
WM 1361, WM 902B, WM 902B, WM 1575, WM 278, WM 1552, WM 35, WM 793: MCDB153 (80%), Leibovitz's L-15 (20%), 2% (v/v) FBS, Insulin 5 μg/ml, $CaCl_2$ 1.68 mM.
IPC298, MEL HO: RPMI 1640, 10% (v/v) FBS, 2 mM Gln

Metastatic melanoma cell lines were cultured as indicated below.
A375, IGR 37 M, IGR-1, CHL1, RPMI 7951: DMEM, 10% (v/v) FBS, 2 mM Gln.
WM 266 M: EMEM, 0.1mM NEAA, 1 mM NaP, 10% (v/v) FBS, 2 mM Gln.
ADMA, SIBR 1949, GALA 1949, CACI, LiGh 1927 B, ElSe 1934, LuPiCi 1936, IrAv 1938, AnSe 1935, SoCh 1970 A: RPMI 1640, 10% (v/v) FBS, 2 mM Gln.
SK MEL 28: DMEM, 10% (v/v) FBS, 0.1 mM NEAA, 2 mM Gln.
C32: EMEM, 0.1mM NEAA, 1 mM NaP, 10% (v/v) FBS, 2 mM Gln.
SK MEL 3: Mc Coy's, 10% (v/v) FBS, 2mM Gln.
G361: EMEM, 0.1mM NEAA, 10% (v/v) FBS, 2 mM Gln.
SK MEL 31: EMEM, 0.1mM NEAA, 1 mM NaP, 10% (v/v) FBS, 2 mM Gln.

MEWO: EMEM, 0.1mM NEAA, 10% (v/v) FBS, 2 mM Gln.
SK MEL 30, COLO 849, COLO 858, COLO 679, COLO 783 and COLO 853: RPMI 1640, 10%(v/v) FBS, 2 mM Gln.

**Western blot analysis**

For protein extraction cells were incubated 30 min on ice in lysis buffer (20mM Hepes pH 7.5, 500mM NaCl, 5mM EDTA, 10% Glycerol, 1% Triton X) and sonicated for 20-40 seconds. 20-50 µg of proteins were separated by 6-10% SDS polyacrylamide gel and transferred to nitrocellulose membranes (Whatman). Immunoblots were probed with the following antibodies: p53 (Santa Cruz DO-1 sc-126), β-catenin (BD Transduction Laboratories, #610154), Vinculin (SIGMA, hVIN-1) and Actin (SIGMA, A240).

**Real-time quantitative PCR (RT-QPCR)**

Total RNA was isolated from cells using RNAeasy (Qiagen) according to the manufacturer's instructions, and treated with DNAse before reverse transcription). cDNA was generated using the M-MuLV Reverse Transcriptase (Finnzymes). The cDNA was used as template in real-time quantitative PCR reactions with Norrin, Axin2, β-catenin specific primers on an Applied Biosystems ABI Prism 7300 Sequence Detection System. The reactions were prepared using SyBR Green reaction mix from Applied Biosystems. RPPo was used as a control gene for normalization.

**Primer sequences for real-time quantitative PCR**

RPPO-fw (TTCATTGTGGGAGCAGAC)
RPPO-rev (CAGCAGTTTCTCCAGAGC)
NORRIN-fw (CTGCGTTCCCCTAAGCTGTG)
NORRIN-rev (TGAAGCTTTCTGGTTGTCATTGTC)
P53-fw (GCTGCTCAGATAGCGATGGTCT)
P53-rev (CATCCAAATACTCCACACGCAA)
CTNNB1-fw (CCCACTGGCCTCTGATAAAGG)
CTNNB1-rev (ACGCAAAGGTGCATGATTTG)
AXIN2-fw (CTCCCCACCTTGAATGAAGA)
AXIN2-rev (GTTTCCGTGGACCTCACACT)

**shRNA sequences**

The targeting sequences of the shRNAs present in the retroviral plasmids are:

shNORRIN#1:
GATCCCCGTGTAGCTCAAAGATGGTGTTCAAGAGACACCATCTTTGAGCTACACTTTTTGGAAA

shNORRIN#2:
GATCCCCACATGTACTAGCTGCATCCTTCAAGAGAGGATGCAGCTAGTACATGTTTTTTGGAAA

GIPZ NORRIN084:
TGCTGTTGACAGTGAGCGCGGATATGTTTAGCTACGTTTATAGTGAAGCCACAGATGTA TAAACGTAGCTAAACATATCCATGCCTACTGCCTCGGA

GIPZ NORRIN086:
TGCTGTTGACAGTGAGCGCGGCACCACTATGTGGATTCTATAGTGAAGCCACAGATGTA TAGAATCCACATAGTGGTGCCTTGCCTACTGCCTCGGA

GIPZ β-CATENIN 022:
TGCTGTTGACAGTGAGCGCGCTCCTTCTCTGAGTGGTAAATAGTGAAGCCACAGATGTATTTACCACTCAGAGAAGGAGCTTGCCTACTGCCTCGGA

GIPZ β - CATENIN 023:
TGCTGTTGACAGTGAGCGCGCTGATATTGATGGACAGTATTAGTGAAGCCACAGATGTAATACTGTCCATCAATATCAGCTTGCCTACTGCCTCGGA

LMP p53:
CCGGCGCACAGAGGAAGAGAAT

**Sh RNA constructs**

The LMP p53 short hairpin RNA construct was generated in the MSCV/LTR/MiR30-PIG (LMP vector) [1]. Oligonucleotides were designed using the "RNAi retriever" (http://katahdin.cshl.org:9331/homepage/siRNA/RNAi.cgi?type=shRNA), and cloned according to published procedures [2].

The shRNA targets a 22 bp sequence at position 1093 in the p53 cDNA, and is reported in the shRNA sequences paragraph above.

**Data mining**

Gene expression data on Norrin and FZD4 were retrieved from the Oncomine website (www.oncomine.org). Data from several studies are shown. Additional details of the studies are available at Oncomine. The study from Miller et al [3] includes 251 primary breast tumours classified according Elston-Ellis histologic grade in low, medium, and high-grade tumours. Expression profiling was performed using the Human Genome U133A and B Array (Affymetrix). The study from van't Veer et al [4] includes 117 breast tumours classified for the histologic grade. Gene expression profiling was performed by using oligonucleotide microarrays containing 24,479 different probes. The study form Bittner et al (International Genomics Consortium, Phoenix, AZ 85004, Expression Project for Oncology - Prostate Sample https://expo.intgen.org/expo/public/ 2005/01/15) includes 60 prostate carcinoma samples analyzed on Affymetrix U133 Plus 2.0 microarrays. Sample data includes grade, stage, Gleason score, and others. This dataset consists of a subset of samples from the Bittner_Multi-cancer dataset.

The study from Bittner et al (https://expo.intgen.org/expo/public/ 2005/01/15; Expression Project for Oncology - Endometrium Samples; International Genomics Consortium, Phoeniz, AZ 85004) includes one hundred seventy-seven (177) endometrial carcinoma samples analyzed on Affymetrix U133 Plus 2.0 microarrays. Sample data includes grade, stage, TNM staging, and others. This dataset consists of a subset of samples from the Bittner_Multi-cancer dataset.

The study from Welsh et al [5] includes 4 normal samples of ovarian tissue and 28 serous papillary ovarian adenocarcinomas. Gene expression profiling was performed by using cDNA microarrays containing more than 6,000 different human genes.

The study from Hendrix [6] includes 103 ovarian samples, comprising 99 tumor samples (37 endometrioid, 41 serous, 13 mucinous, and 8 clear cell carcinomas), and 4 normal tissue samples, which were analyzed on an Affymetrix HG-U133A array.

The study from Dhanasekaran et al [7] includes samples from normal prostate adjacent to tumors (2), normal adult prostate (7), normal pubertal prostate (3) and prostate tumors (25). Gene expression profiling was performed by using Microarrays (20K chip) containing sequence – verified, PCR-amplified human cDNAs representing 15,495 Unigene clusters. The study from Welsh et al [8] includes normal prostate tissues (9) and prostate tumors (25);

eight of the cancers were paired with normal tissue obtained from the same patient. Gene expression profiling was performed by using the Human Genome U95A-Av2 Array (Affymetrix), representing ~8920 different genes.).

**Statistical analysis**

A two-tailed, unpaired Student's t test was done to determine the statistical significance by the probability of difference between the means. $P<0.05$ is considered statistically significant.

**Legends to Supplementary Figures**

**Supplementary Figure 1. Inhibition of Norrin expression by different RNA interference constructs.**

**a.** pRetroSUPER – based Norrin sh#1 inhibits Norrin expression. RT-QPCR analysis of Norrin mRNA. The mRNA was prepared from the experiment shown in Figure 1d. Expression of Norrin mRNA is expressed relative to RPPo. Error bars represent the mean +/- SD for a representative experiment performed in triplicate.

**b.** Inhibition of Norrin expression by the GIPZ Norrin 086 lentiviral construct (Open Biosystems). RT-QPCR with Norrin specific primers. The mRNA was prepared from the experiment shown in Figure 1e.

**Supplementary Figure 2. Norrin activates the WNT/β-catenin pathway.**

**a, b.** Inhibition of Norrin expression by two lentiviral RNA interference constructs (GIPZ Norrin 084 and GIPZ Norrin 086) targeting different regions of Norrin, leading to downregulation of the WNT/β-catenin pathway target gene *AXIN2*. RT-QPCR with primers specific for AXIN2, or Norrin, using mRNA from cells infected with the indicated constructs.

**c.** rhNorrin exposure upregulates the WNT/β-catenin target gene *AXIN2*. RT-QPCR for AXIN2, in TIG3-T cells treated with rhNorrin (60 ng/ml) for 12 hours.

**d.** rhNorrin increases the levels of the β-catenin protein. Immunoblot analysis of U2OS cell extracts, showing β-catenin levels and Vinculin as a loading control. From the left lane: non treated control, cells treated for 24 hours with conditioned medium from L1 cells expressing or not WNT3a (1:3 dilution), with recombinant human Norrin (20ng/ml or 40 ng/ml), or 2 mM LiCl.

**Supplementary Figure 3. Recombinant Norrin inhibits cell proliferation dependent on p53.**

**a.** TIG3-T cells non-treated or treated with rhNorrin (40ng/ml), conditioned medium from L1 cells (1:3 dilution) expressing secreted WNT3a, or conditioned medium from L1 cells without the expression of WNT3a. Cumulative increase in cell number is shown versus time in days.

**b.** Growth curves of TIG3-T cells in the absence or presence of rhNorrin (80ng/ml).

**c.** Growth curve of TIG3-T cells infected with a lentiviral plasmid expressing shRNA to p53 (LMPp53) in the presence or absence of rhNorrin (80ng/ml).

**d.** Immunoblot analysis of extracts from cells in panel c, showing p53 protein levels.

**Supplementary Figure 4. Norrin expression is significantly decreased in several human tumours.**

**a.** Norrin expression data from two microarray expression studies in ovarian cancers, retrieved from the Oncomine database. P-values of T-tests are indicated.

**b.** Norrin expression in two studies of prostate primary tumours. Data plot from Oncomine database. P-values of T-tests are indicated.
