## Supplementary Figures for "The Frizzled-ligand Norrin acts as a tumour suppressor linking oncogenic RAS signalling to p53"

**a**

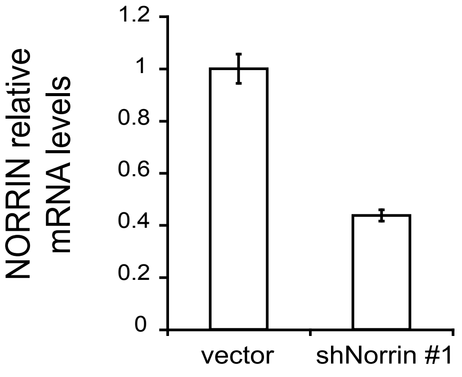


**b**

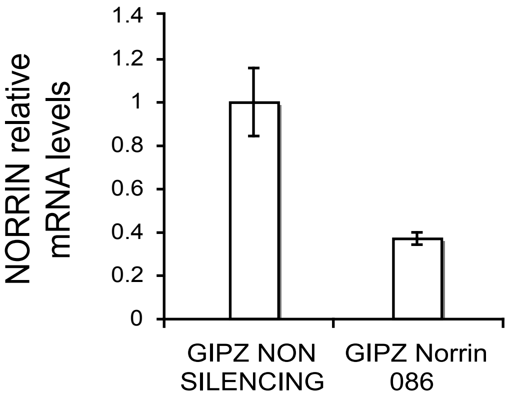


**Supplementary Figure S1**

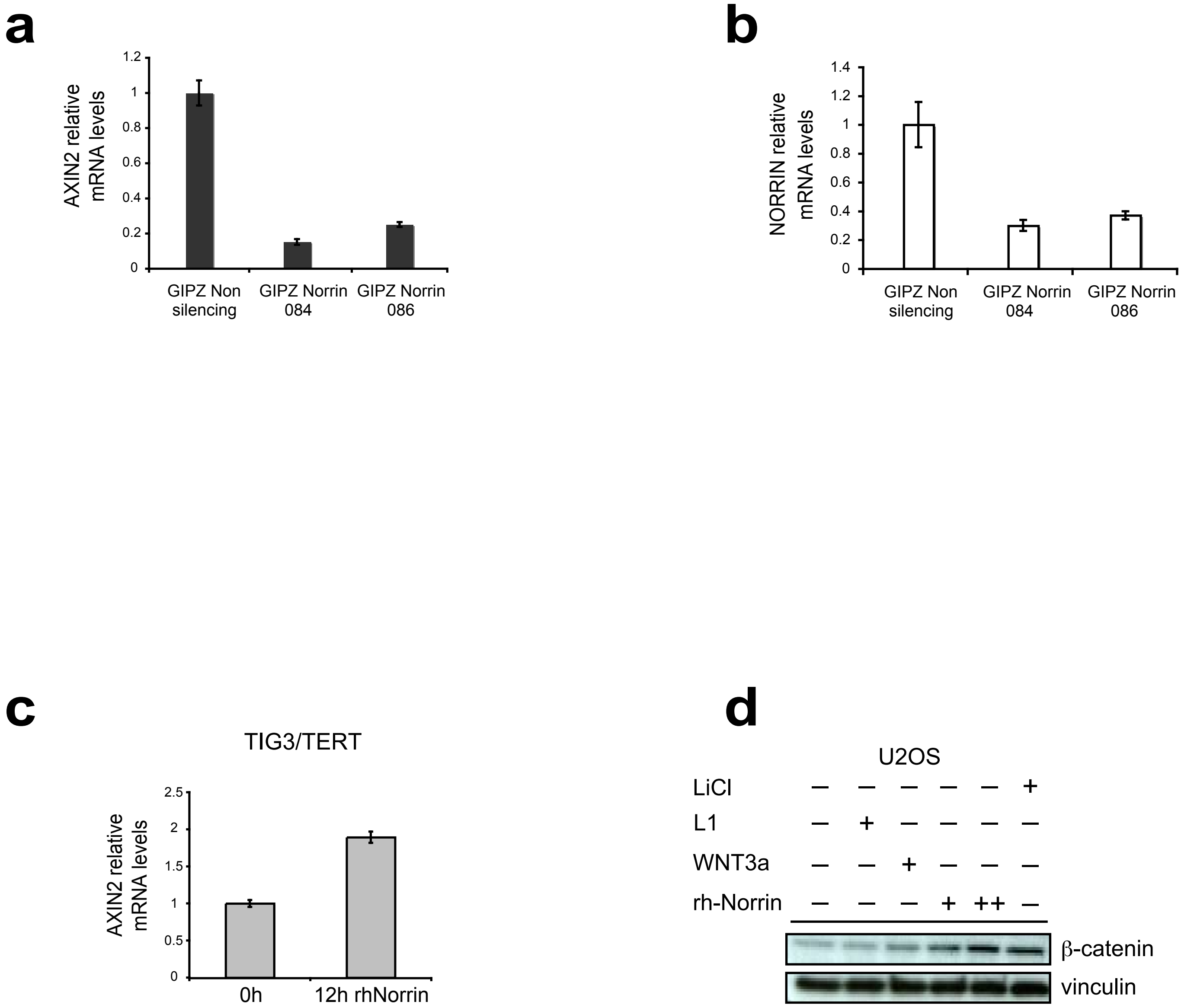


**Supplementary Figure S2**

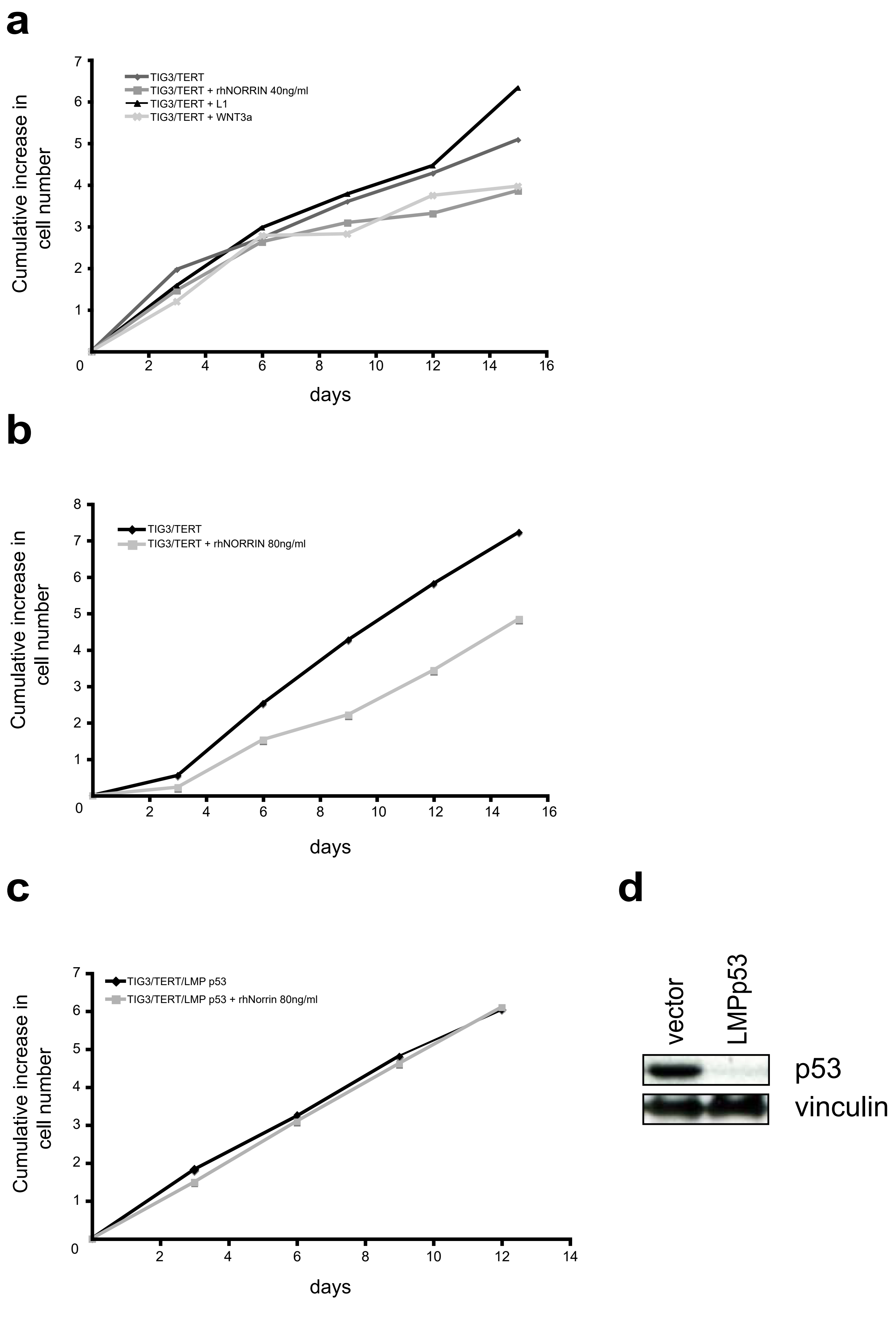


**Supplementary Figure S3**

**a**

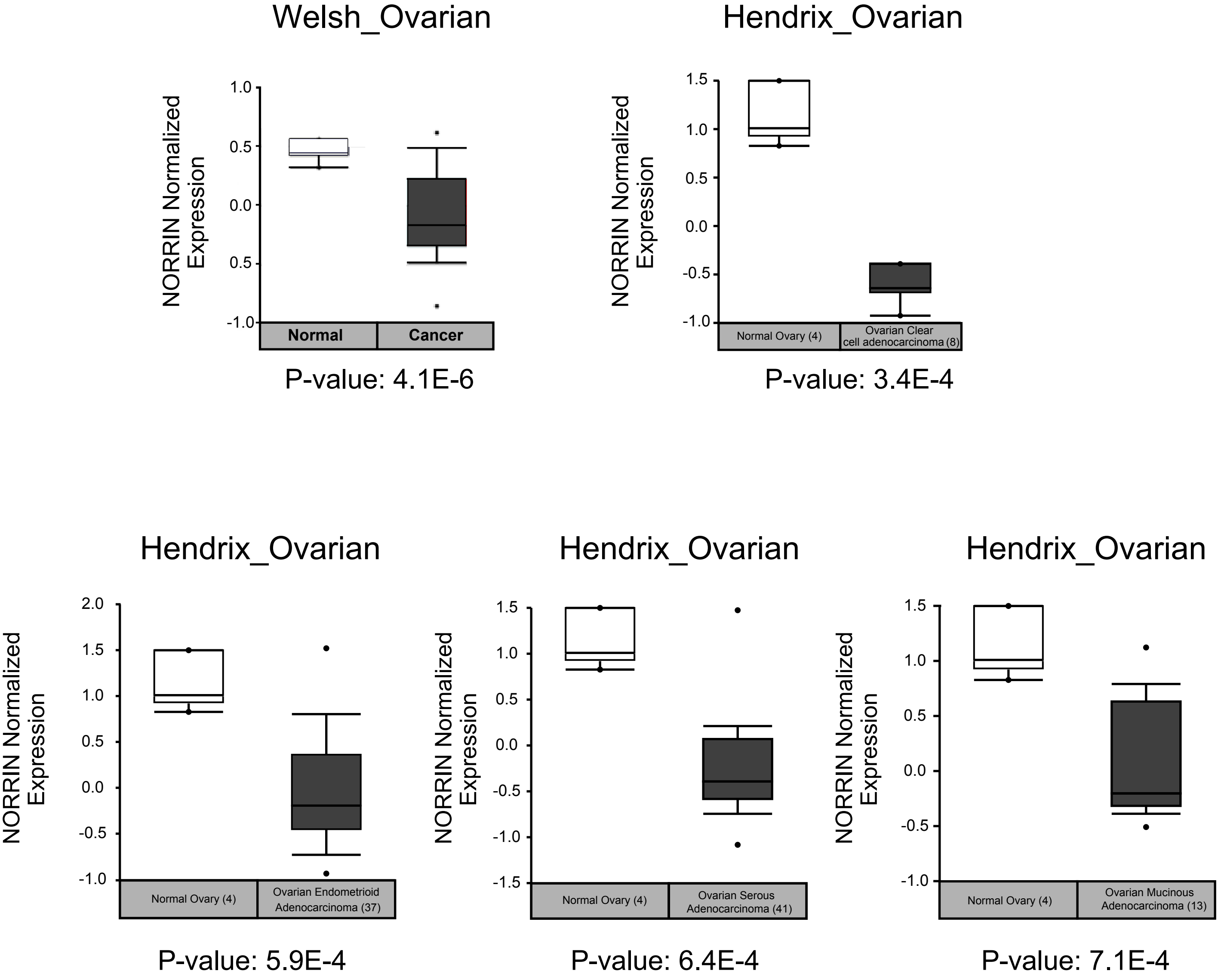


**b**

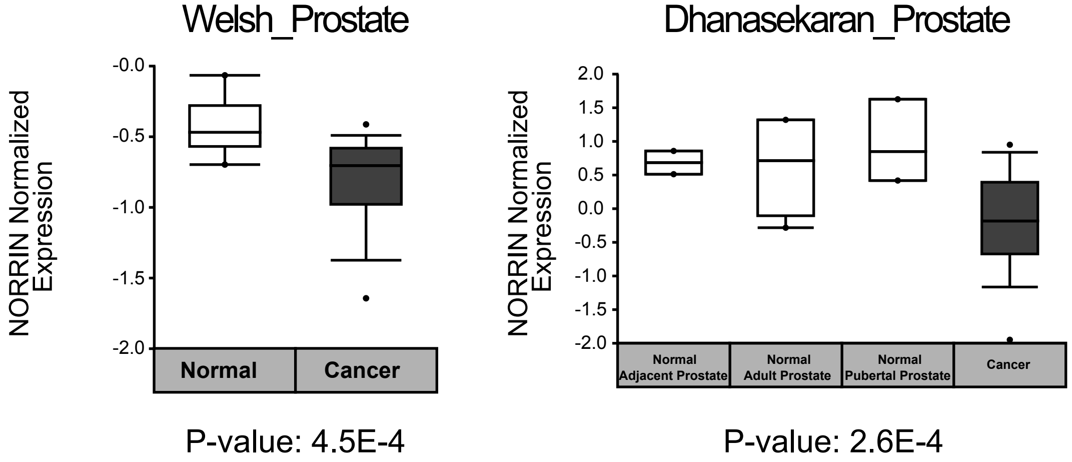


**Supplementary Figure S4**
